# Factors that influence scientific productivity from different countries: A causal approach through multiple regression using panel data

**DOI:** 10.1101/558254

**Authors:** Bárbara S. Lancho-Barrantes, Héctor G. Ceballos, Francisco J. Cantú-Ortiz

## Abstract

The main purpose of the economic expenditure of countries in research and development is to achieve higher levels of scientific findings within research ecosystems, which in turn could generate better living standards for society. Therefore, the collection of scientific production constitutes a faithful image of the capacity, trajectory and scientific depth assignable to each country. The intention of this article is to contribute to the understanding of the factors that certainly influence in the scientific production and how could be improved. In order to achieve this challenge, we select a sample of 19 countries considered partners in science and technology. On the one hand we download social and economic variables (gross domestic expenditure on R&D (GERD) as a percentage of gross domestic product (GDP) and researchers in full-time equivalent (FTE)) and on the other hand variables related to scientific results (total scientific production, scientific production by subject areas and by different institutions, without overlook the citations received as an impact measure) all this data within a 17-year time window. Through a causal model with multiple linear regression using panel data, the experiment confirms that two independent (or explanatory) variables of five selected explain the amount of scientific production by 98% for the countries analyzed. An important conclusion that we highlight stays the importance of checking for compliance of statistical assumptions when using multiple regression in research studies. As a result, we built a reliable predictive model to analyze scenarios in which the increase in any of the independent variables causes a positive effect on scientific production. This model allows decision maker to make comparison among countries and helps in the formulation of future plans on national scientific policies.

## 1 Introduction

Research is an ensemble of activities performed by academicians to produce new knowledge contributing with this to the development and progress of entire society. Research results mainly are disseminated through scientific papers, reviews, conference proceedings, even monographs or book-chapters etc. [1]. The resulting collection of documents is known as scientific production which may correspond to the productivity of a year, a specific period, a researcher, a research group, an institution, a discipline, or even a whole country [2].

The amount of scientific production can increase, decrease or become stagnant because is an unfixed parameter influenced by many other external features of economic, sociological, cultural, and political nature [3]. In fact the scientific literature encompasses diverse works that demonstrate that scientific production is influenced by institutional conditions, cultural dimensions, knowledge management processes, barriers in access to information, technological and human capital, such as databases, scientific resources, software licenses, well-equipped laboratories, material in optimum conditions, and personnel dedicated to research & development (R&D), etc. [1, 4–6].

Barjak [7] classified factors that affect scientific production into individual personal circumstances, such as research motivation, experience, personality, mobility and adaptation to change, creativity, age, gender, professional range, recognition, compulsory teaching, administrative and management assignments, communication skills with colleagues, and participation in international research networks. In addition, scientific production could be influenced by environmental and sociological conditions, such as idiosyncrasy of countries, demographic traits, disciplines patterns, institutional thematic preferences, research group sizes, institutional prestige, and research freedom.

Collaborative factors have also been considered as influential. In addition, new information technologies allow researchers collaborate in different geographic areas to share resources, skills, and competencies. Furthermore, many works have manifested that high levels of scientific collaboration lead to greater research productivity, greater papers quality, and higher levels of impact and citations in publications [8–15].

Some studies have revealed a significant relationship between scientific production and the global economy [16–19]. Other studies have described the relationship between inputs (i.e., funding and economic investments in R&D) and outputs (i.e., generated results) [1, 20–23]. Other studies have analyzed the impact of gross domestic product (GDP) and investment in research on the number of universities and indexed journals and citations in different countries [24–27].

However, other studies have shown that simple correlation analysis could not reveal relation among variables because could not be entirely accurate to infer real causality with such kind of analyses [18, 19]. Lee, Lin, Chuang, & Lee [28] and Ntuli, Inglesi-Lotz, Chang, & Pouris [29] examined the causal relationship between research production and economic productivity applying the bootstrap panel Granger causality, and Inglesi-Lotz et al. [17] also applied panel causality test to examine the causal relationship between research performance and economic growth using Brazil, Russia, India, China, and South Africa (BRICS) data.

Furthermore, there is a large number of theoretical works which have demonstrated the relationship of these variables with scientific production: economic investment in science [20, 29], researchers [22, 30], institutions [24–26, 31], disciplines [32–34], and the citations obtained [35, 36].

Nevertheless, to date, no single criterion could identify the influencing determinants because scientific production could be altered by other imperceptible external factors. In order to contribute to the collection of studies concerned about the behaviour of scientific production we have elaborated a regression model using panel data to examine the scientific production of strategic countries through a succession of influential variables at the same time.

We selected a sample of 19 countries considered partners in science and technology with data from 17 year time window. The originality and main advantage of this study is to apply multiple regression using panel data to a diverse set of countries considered strategic partners in science and technology. Through panel data we can capture unobservable heterogeneity, either between economic agents or studies over time, because heterogeneity cannot be detected by time-series or cross-sectional studies. Multiple regression with panel data enables more dynamic analysis by incorporating the temporal dimension, which enriches the study, particularly in periods of significant and multiple changes. Panel data models are frequently used in statistic and econometric studies. Multiple regression enables analysis of two important aspects when working with panel data that form part of the unobservable heterogeneity, i.e., specific individuals and temporary effects. This technique allows researchers to have a greater number of observations, improving of information quality and efficiency because increasing the sample size, we obtain more information about the population and, consequently, the degrees of freedom increase [37].

In order to choose the explanatory variables, we followed an empirical procedure based on previous statistical experiments with scientific production variable. Beforehand we selected a large number of variables which we consider influencing scientific production however they were ultimately reduced by statistical procedures. Note that even without an empirical procedures we could have affirmed that some external variables can influence scientific production by research logic and observation of this phenomenon.

The variables chosen are the following: the most important is expenditure in research, without this economic input we could not do anything in science. The second one is the human labor ‘researchers’, is also something reasonable, since they represent the human force to perform research production. Moreover, the third is the countries’ research preference measured by scientific production in disciplines. The fourth variable is the higher education and research institutions because they are responsible for hosting the scientific processes playing an enormous importance in the production. Finally, citations were selected as another exogenous variable. Citations motivate researchers to continue producing, collaborating, or developing a specific productive research line- when a group of researchers collaborate and these collaborations are successful with respect to the impact received and citations, this motives to continue collaborating. Consequently citations is a variable that encourages collaboration and with collaboration increases the scientific production.

Therefore, we have chosen the variables which consider the most relevant for our study. We believe that could make governments reconsider their scientific policies when assigning their economic resources.

A number of research questions are formulated: Could the selected variables together explain the scientific production behaviour? If governments increase investment in science, by 1%, would scientific production increase? Would increasing the number of researchers improve scientific production? Are a small group of institutions responsible for increasing scientific production? Is a concentrated number of disciplines responsible for increased scientific production? Does the total number of citations motivate researchers to continue publishing? Our working hypothesis is as follows: variations in scientific production can be explained by the previously identified descriptive variables (economic expenditure, researchers, research preference, academic and research institutions, citation received) considering that these variables show dynamic behaviour over time. In other words, we contend that a causal relationship between scientific production and exogenous variables can be captured by regression analysis using panel data.

We are aware that investment in research is not allocated in an equal manner in all scientific areas, however we have analyzed all countries with the same variables to make a comparison. In a future study we could dismember the economic investment destined to the different areas and also account for the resulting production in those areas. Similarly we would like to point out that investment in research is channeled and destined to different sectors, however we consider that production can be seen as a way to materialize that economic injection.

In order to select a sample of countries we choose Mexico and its 18 strategic countries in science and technology [38]. This is a diverse, consolidated, defined, closed, easily controllable and heterogeneous sample which could be an example closer to the behaviour of the world reality. For this we consider this as an adequate sample to answer our questions and test the hypothesis since this conforms a wide sample of countries.

Some bibliometrics studies focusing on Mexico have been published. The most recent is about the general scientific production of Mexico [39]. Moreover Arvantis, Russel, Rosas, & A., [40] analyzed articles from Mexico in the National Citation Report database. Lima, Liberman, & Russell, [41] studied the relationship between the number of links among an article’s authors and a Likert scale-based measure of group cohesion in three research areas, i.e., biotechnology, mathematics, and physics, at the National Autonomous University of Mexico (UNAM). Castillo-Pérez, Muñoz-Valera, García-Gómez, & Mejía-Aranguré [42] analyzed the volume and impact of Mexican scientific production relative to influenza published in the Science Citation Index between 2000 and 2012. Uddin, Singh, Pinto, & Olmos [43] published a detailed, text-based, scientometric analysis of computer science research in Mexico from 1989 to 2014 indexed in the Web of Science. Franco-Paredes, Díaz-Reséndiz, Pineda-Lozano, & Hidalgo Rasmussen, [44] characterized the scientific production of the Mexican Journal of Eating Disorders during the period of 2010–2014. Frixione, Ruiz-Zamarripa & Hernández [45] conducted a limited survey of the functioning and results of the 30+-year-old National System of Researchers, which is Mexico’s primary instrument to stimulate competitive science and technology research. Hernandez-Garcia, Chamizo, Kleiche-Dray, & Russell [46] studied steroid research between 1935 and 1965. The bibliometrics and searches in patent files in their paper indicated that the Syntex industrial laboratory in Mexico and the National Autonomous University of Mexico (UNAM) produced approximately 54% of the documents in the main journals, which in turn generated more than 80% of the citations in this period. Villaseñor, Arencibia, & Carrillo-Calvet [47] produced multiparametric scientometric characterizations of the production profiles of the 50 most productive Mexican higher education institutions listed in Scopus, Elsevier’s international bibliographic database [48]

## 2 Material and methods

In order to perform this study, we chose a sample of strategic partners’ countries which are different in cultural, social, and economic magnitudes and there is a representation of all regions of the world. The countries’ sample is identified in Conacyt’s latest general report on science, technology, and innovation [38], as shown in Table 1.

**Table 1.** Strategic countries [38]

| Region | Countries |
| --- | --- |
| Africa | South Africa |
| Asiatic Region | China, Japan, South Korea, and India |
| Eastern Europe | Russian Federation |
| Latin America | Argentina, Brazil, Colombia, Chile, Mexico |
| Middle East | Israel and Turkey |
| Northern America | Canada and the United States |
| Western Europe | Germany, Spain, France, and the United Kingdom |

**Table 2.** Non-standardized estimates of the general model with variables GERD/GDP and A&RI without country particularity.

| | $\beta$ | Sig. |
| --- | --- | --- |
| $\beta_0 =$ (Constant) | 7.395 | 0.000 |
| $\beta_1 =$ GERD/GDP | 0.723 | 0.000 |
| $\beta_2 =$ A&RI | 1.169 | 0.000 |

We used Scopus to extract scientific variables (scientific production, scientific production by disciplines and institutions) because it represents the overall structure of world science at a global scale. Scopus is the world’s largest scientific database which is the most comprehensive and accurate bibliometric database widely used in diverse studies. It covers journals included in the Thomson Scientific Web of Science (WoS) and more [49], and its coverage is statistically balanced in terms of subjects, countries, languages, and publishers. We have also used SciVal to extract the total number of citations received.

For the economic component of the study, we used statistics provided by the United Nations Educational, Scientific, and Cultural Organization (UNESCO) Institute for Statistics (UIS) [50] to extract the gross domestic expenditure on research and development (GERD) as a percentage of GDP (GERD/GDP) as well as number of researchers per million inhabitants calculated as a full-time equivalent (FTE).

UNESCO takes the data from the OECD database. However, it is worth pointing out that many of strategic countries do not belong to the OECD; therefore, to give consistency to the sources, we decided to use only the data from UNESCO statistical database instead of OECD’s data.

The timeframe for the economic part of our study (GERD/GDP and Researchers) was 1996–2012. The oldest data are from 1996, which is the first year for which complete country data exists in the UNESCO database. To measure the effect of investment in research on scientific productivity, we displaced the scientometric data by three years. The three-year displacement period was employed because the effect of GERD/GDP investment on scientific production takes at least three years to manifest. We determined the three-year displacement period by experimenting with several time windows.

The time frame for the temporal data in the scientometric sample (i.e., scientific production, citations, institutions, and disciplines) was 1999–2015. The temporal data, both economic and scientometric, encompass a total of 17 years.

We downloaded data from Scopus, SciVal, and UNESCO in March 10, 2017.

Variable definitions:

### Dependent or outcome variable

#### Scientific production

Total number of documents produced by the target countries (1999–2015). All document typologies are considered. Note that in order to assign the same weight to all countries we used whole counting instead of fractionalized ones to measure research output of countries.

### Independent or predictor variables

#### GERD as a percentage of GDP

Gross domestic spending on R&D is defined as the total expenditure (current and capital) on R&D by all resident companies, research institutes, university and government laboratories, etc., in a country. It includes R&D funded from abroad but excludes domestic funds for R&D performed outside the domestic economy. This indicator is measured in million USD and as a percentage of GDP [50]. Note that in this context, GDP is defined as the sum of the gross value contributed by all resident producers in the economy, including distributive trades and transport. Here GDP includes product taxes but does not include subsidies, i.e., subsidies not included in the value of the products have been subtracted. This measure is defined to better understand GERD/GDP; however, it has not been taken as an independent variable [50].

We used GERD, as this indicator groups the investment data in a global way (both government, business, other organizations, etc.). This refers to investment in research and technological development. GERD expenditure broken down into other four indicators like HERD - Higher Education Expenditure on R&D, GOVERD - Government Expenditure on R&D, BERD - Business Enterprise Expenditure on R&D, and Private Non-Profit Research and Development (PNP).

#### Researchers (FTE)

Represents the number of professionals engaged in the conception or creation of new knowledge, i.e., professionals who conduct research and improve or develop concepts, theories, models, instrumentation techniques, software, or operational methods, during a given year expressed as a proportion of a population of one million. Researchers FTE is calculated as the number of researchers during a given year divided by the total population (using the mid-year population as reference) multiplied by 1,000,000 [50].

#### Academic & research institutions (A&RI)

Total number of institutions responsible for at least 50% of the production of each country in a year. We identified only institutions that generated at least 50% of the annual scientific production during the study period. We used this information to determine institutions’ influence in scientific production.

#### Subject areas

The total number of subject areas involved in at least 50% of the production of each country in a year. We determined how many of the 27 subject areas included in Scopus are included in at least 50% of the scientific production to verify if the degree of concentration or dispersion in research disciplines influences the behavior of general scientific production.

#### Citations received

Total number of citations (up to last data cut) for documents published in a year.

### Multiple linear regression using panel data

Multiple linear regression at a significance level of 5% (α = 0.05) was used to determine causality between scientific production and the predictor variables. Here the dependent variable (Y) represents scientific production and the explanatory and independent variables (X) are GERD/GDP, researchers, A&RI, subject areas, and citations. Since our sample combines temporal and transversal dimensions, the most adequate model to explain causality can be obtained using panel data. This will allow us to analyze the general effect and observe the individual outcome for each country in consideration of the influence of explanatory variables on the dependent variable.

To measure this effect, we have also created binary dummy variables to quantify the effect of the country on the scientific production variable, which takes a value of 1 when the analyzed country is present and 0 when the country is not present.

When extracting data from the UNESCO database, we detected incomplete data for some countries (i.e., Israel, South Africa, India, Colombia, Turkey, Brazil, and Chile); therefore, to maintain accuracy we decided not select these countries in the present study. An analysis of factors influencing their scientific production could be explained with other methods in future works.

As a result, a total of 12 countries with 204 observations for the 17-year period were considered in this study.

### 2.1 Statistical assumptions

We consider that checking the validity of statistical assumptions required by multiple regression using panel data is a particular strength in this study. We deliberately decided to mention in this paper the variables that did not comply with the statistical assumptions instead of removing them or choosing others. We noticed that in some published studies where regression analysis is used, statistical assumptions are not mentioned, obviated, or only validated with respect to the multicollinearity assumption. This could lead to unreliable and imprecise results and this is something we avoid in this study.

We consciously leave the variables that did not meet the assumptions to alert the scientific community of an incomplete use of the regression in bibliometrics could damage the scientific results.

We performed our statistical analyses using the SPSS (version 24) statistical package. We tested the statistical assumptions of the multiple linear regression modeling because, to create inferences about Y from the sample data, it is necessary to establish assumptions about the behavior of error ε, which defines the random behavior of Y, and perform experiments according to these assumptions.

The assumptions of the multiple regression modeling we evaluated are as follows.

#### Linearity assumption

The regression model assumes that the relationship between the dependent and independent variables is linear; however, in practice, some variables demonstrate curvilinear (i.e., nonlinear) relationships. Note that estimating a linear regression model with variables that have nonlinear relationships results in unreliable and imprecise estimates.

#### Multicollinearity assumption

In addition to linearity, another principle of regression modeling is that the explanatory variables should not be correlated with each other. When two explanatory variables are strongly correlated, a collinearity problem exists, and when more than two are correlated, we have a multicollinearity problem. We used the following to identify if such problems were present: a matrix of correlations between explanatory variables, the variance inflation factor (VIF), multicollinearity diagnoses, and the proportion of variance. As per the correlation matrix, if two or more variables have a correlation coefficient greater than or equal to 0.9, there is a collinearity or multicollinearity problem. If the VIF is greater than or equal to 10, there are collinearity or multicollinearity problems. With the multicollinearity diagnoses, we can check the condition index, which measures the association between independent variables. Its value is the square root between the largest and the smallest eigenvalue. If its value is greater than or equal to 30, there are strong multicollinearity problems as long as this value is attributed to the explanatory variables. The proportion of variance measures the origin of multicollinearity. It represents the proportion of the variance that each eigenvalue has in each explanatory variable. If two or more variables have a ratio of 0.9 or greater, this indicates that those variables have a multicollinearity problem.

#### Assumption of normality

For any combination of the values of X, variable Y must have normal distribution. Failure to comply with this assumption invalidates the statistical tests performed on the regression coefficients and the future values of Y. In our case, this assumption is the easiest to validate given that we have a large sample (n≥30), and in practice, according to the Central Limit Theorem of large samples, we conclude that our data meet the normality assumption.

#### Extreme and influential observations assumption

An observation that is distant from the rest of the data is considered an outlier observation. Both extreme and influential observations affect estimations because they considerably modify estimates, i.e., standard errors of high coefficients, low determination coefficients, and coefficients with signs or with magnitudes that are significantly different from their true values. We evaluated this using the Cook Distance criterion. An observation can be considered influential if the Cook Distance is greater than or equal to 1.

#### Assumption of independence

There should be either no dependence or correlation between the values of the error term ε and between the values of the “y” variable. Violation of this assumption is known as autocorrelation. One corrective measure to assure the assumption of independence is the Cochrane–Orcutt transformation. However, we used the Durbin–Watson test to validate if the independence assumption is satisfied. The value of the test statistic “d” ranges from 0 to –4, where small values close to 0 indicate a positive autocorrelation and large values indicate a negative autocorrelation.

Additionally, we used the Newey–West estimators during the regression to try to overcome heteroscedasticity introduced by the differences among national research policies. Despite the use of these estimators do not change the value of the coefficients obtained without the Newey-West correction, it corrects their significance in some cases.

Once we checked the statistical assumptions for this study, we detected noncompliance with some of the selected variables:

Researchers (FTE): This variable presented strong collinearity with GERD/GDP. This appears somewhat logical because researchers are a consequence of investment in science. We decided to keep GERD/GDP in the regression because this is an essential variable controlled by the government.

Subject areas: When data were extracted from the Scopus database, we found that this variable remained constant for all countries over time, and as is well known, regression requires variability in data, particularly when making forecasts [51]. The number of subject areas with almost 50% of scientific production in all countries ranged from three to six in each year.

Citations: Linearity is the first requirement of multiple regression. This variable demonstrated a curvilinear form for all countries, which was impossible to correct by any type of linearity transformation.

Finally, we obtained two explanatory variables: GERD/GDP and A&RI in addition to the dummy variables of the countries. With these variables, we validated and fulfilled the five statistical assumptions: linearity, multicollinearity, extreme and influential observations, normality, independence and also heteroscedasticity. With multiple linear regression using SPSS, we obtained a model comprising the following two equations.

The first equation includes the dependent variable (Y) scientific production, and the independent variables (Xs), GERD/GDP (X_1_) and A&RI (X_2_), as well as the corresponding dummy variables for each country. For a better estimation of the data, all variables, including the dependent variable, were transformed by applying a natural logarithm, except for the dummy variables because they are binary variables. If the effect of the country (dummy variables) is not present, we obtain the following equation.

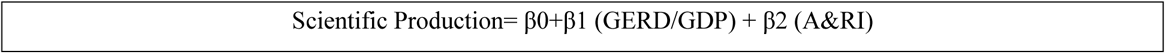

The second equation considers that the 12 countries are present, and the estimated model allows us to analyze the particularity of each country. This equation is expressed as follows.

Scientific Production = β0 + β1 (GERD/GDP=X_1_) + β2 (A&I=X_2_) + β3 (Dummy Argentina) + β4 (X_1_ * Dummy Argentina) + β5 (X_2_ * Dummy Argentina) + β6 (Dummy Canada) + β7 (X_1_* Dummy Canada) + β8 (X_2_* (Dummy Canada) + β9 (Dummy France) + β10 (X_1_* Dummy France) + β11 (X_2_* Dummy France) + β12 (Dummy Germany) + β13 (X_1_* Dummy Germany) + β14 (X_2_* Dummy Germany) + β15 (Dummy Spain) + β16 (X_1_* Dummy Spain) + β17 (X_2_* Dummy Spain) + β18 (Dummy United Kingdom) + β19 (X_1_* Dummy United Kingdom) + β20 (X_2_* Dummy United Kingdom) + β21 (Dummy United States) + β22 (X_1_* Dummy United States) + β23 (X_2_* Dummy United States) + β24 (Dummy China) + β25 (X_1_* Dummy China) + β26 (X_2_* Dummy China) + β26 (X_2_* Dummy China) + β27 (Dummy Japan) + β28 (X_1_* Dummy Japan) + β29 (X_2_* Dummy Japan) + β30 (Dummy South Korea) + β31 (X_1_* Dummy South Korea) + β32 (X_2_* Dummy South Korea) + β33 (Dummy Russia Federation) + β34 (X_1_* Dummy Russian Federation) + β35 (X_2_* Dummy Russian Federation)

In this model, Mexico’s data are used as a reference (intercept). To check the country effect, the country’s value is compared to the reference Mexico values.

Here parameter β0, the “coordinate at the origin,” tells us how much Y increases when all X = 0. Parameter β1, the “slope,” indicates the increase in Y for each increase of 1% to X_1_. The same applies to parameters β2 and X_2_. Examples:

Equation 1 of Mexico: Scientific Production = β0 + β1 X_1_ + β2 X_2_, where all dummy variables equal 0.

Equation 1 of Spain: Scientific Production = β_0_+β_1_ X_1_ + β_2_ X_2_ + Dummy Spain + β3 X_1_Dummy Spain + β4 X_2_ Dummy Spain, where dummy Spain = 1 and the other dummies = 0.

In the Spain equation, X_1_ and X_2_ correspond to GERD/GDP and A&RI, respectively.

We can observe how the panel data model combines in the same equation cross-section data and temporal cut data to demonstrate causality. This model provides more information, more variability, less collinearity among variables and a higher precision. Finally, the data for the very valuable information to individuals following them through time, offers a more complete view of the problem, interpreting the dynamics of the change in cross-sections. One of the most important advantages of panel data with respect to other types of data is that they allow us to control unobservable differences.

We work a random effects model, ε it is assumed to vary stochastically over i or t requiring special treatment of the error variance matrix.

## 3 Results and Discussion

In this section, we show the parameters estimated by multiple linear regression modeling using panel data. We show the general estimates with the predictor variables GERD/GDP and A&RI without the country effect. We also show the estimates with the dummy variables to highlight the presence of the different countries.

The first estimates correspond to the following general equation.

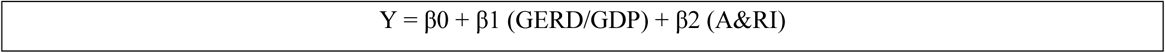

As we can observe, the GERD/GDP and A&RI variables explain the dependent variable. Here the level of significance is 0.000; therefore, the confidence level lies in the 95%–100% range.

Table 3 shows that the adjusted R^2^ value is 0.73, which means that the two predictor variables explain the scientific production variable with 73% accuracy, which is quite acceptable. The predictive value of the model with its two independent variables is high, as shown by the F values and statistical significance.

**Table 3.**
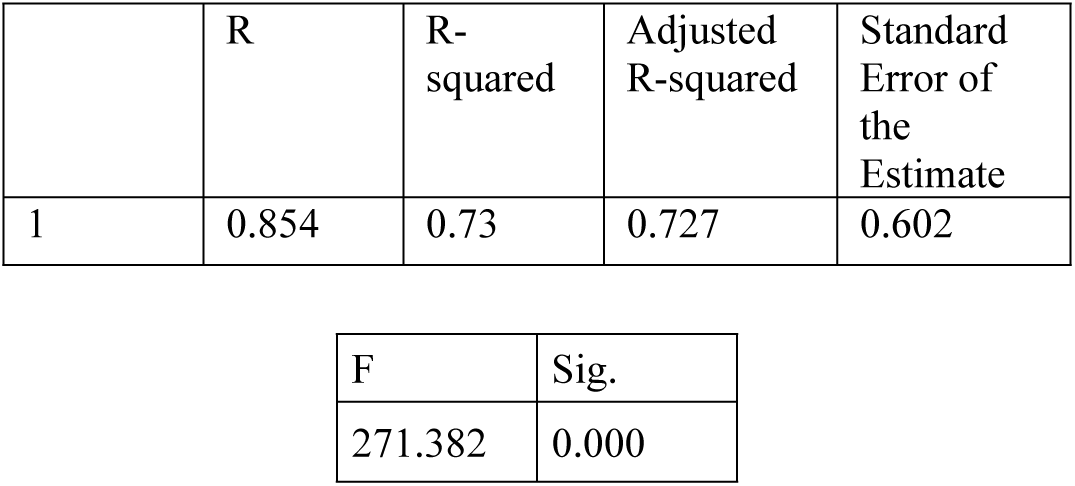
R^2^ adjusted, test F, statistical significance and standard error of the estimate of the general model

When the GERD/GDP and A&RI variables tend to 0 in the analyzed countries, on average, the scientific production is 1628 (e + 7.395). The effects are seen when the percentage changes. For example, when the GERD/GDP variable increases by 1%, the effect on scientific production will be 8.118% provided that the value of the A&RI variable remains constant. In contrast, if the number of institutions increases by 1% and GERD/GDP is constant, the scientific production increases to 8.564%. In general, we observe that both variables have a positive influence on scientific productivity.

Table 4 shows the estimates of the model. This allows us to analyze the particularity of each country by applying the dummy variables.

**Table 4.** Non-standardized coefficients and significance of the model with GERD/GDP and A&RI variables in consideration of country presence

| Country | Intercept/Dummies | | GERD/GDP ($X_1$) | | A&RI ($X_2$) | |
| --- | --- | --- | --- | --- | --- | --- |
|  | Coefficient | Sig. | Coefficient | Sig. | Coefficient | Sig. |
| Mexico | 7.068 | 0.000 | 1.438 | 0.000 | 1.436 | 0.000 |
| Argentina | 7.217 | 0.000 | -1.695 | 0.000 | -4.017 | 0.000 |
| Canada | 18.868 | 0.003 | 0.436 | 0.510 | -7.395 | 0.001 |
| China | -0.020 | 0.989 | 0.128 | 0.676 | 0.113 | 0.817 |
| France | 13.965 | 0.000 | -3.753 | 0.000 | -3.934 | 0.000 |
| Germany | -2.312 | 0.000 | 0.928 | 0.001 |  |  |
| Japan | 3.379 | 0.004 | 0.034 | 0.900 | -1.588 | 0.000 |
| Russian Federation | 1.752 | 0.078 | -1.202 | 0.000 | -0.603 | 0.065 |
| South Korea | -4.981 | 0.000 | -0.946 | 0.000 | 1.184 | 0.000 |
| Spain | 0.107 | 0.918 | 0.297 | 0.180 | -0.286 | 0.378 |
| United Kingdom | 15.055 | 0.000 | -2.403 | 0.326 | -4.519 | 0.000 |
| United States |  |  | 2.782 | 0.085 | -0.913 | 0.032 |

**Table 5.** Adjusted R^2^, test F, statistical significance, and standard error of the estimate of the model in consideration of country particularity.

| Model | R | R-Squared | Adjusted R-Squared | Standard Error of the Estimate |
| --- | --- | --- | --- | --- |
| 1 | 0.995 | 0.990 | 0.988 | 0.125 |

| F | Sig. |
| --- | --- |
| 517.733 | 0.000 |

When we incorporate the country variable, prediction improves compared with the prediction of the previous general model. Here the adjusted R^2^ takes a value of 0.988, which means that these two variables explain the dependent variable for all countries by 98%. Therefore, panel data with dummy variables improve the estimate by reducing the error from 0.602 to 0.125. The standard error of the estimate represented by the letter “S” is used to make inferences about the fit of the data to the regression equation, it is also the point estimate of the standard deviation of the error and Y. Models will be preferred where S is closest to zero.

As shown in Table 4, the model excludes the dummy variable for the United States because the dummy data of this country had the highest collinearity (close to 1) in data for the A&RI variable as the number of institutions that are responsible for 50% of the United States’scientific production remains constant over the years. Another excluded variable is Germany_A&RI because it has high correlation with Germany’s GERD/GDP. However, even when the regression model excludes these variables, their effect will be measured and added to the effect of the rest of the countries in a global manner according to their statistical significance.

The following figures show the different effects of the different countries based on the estimates shown in Table 4.

As can be seen in Table 6, the United States dummy variable is significant with respect to scientific production. However, the dummy variable for Germany’s A&RI is not significant and will therefore have the effect of the reference country, i.e., Mexico.

**Table 6.** Excluded variables

| Model | $\beta$ | Sig. |
| --- | --- | --- |
| Dummy United States | 7.720 | 0.003 |
| Dummy Germany<br>A&RI | 1.071 | 0.376 |

**Table 7.** Percentage of scientific collaboration documents classified by international, national, and institutional collaborations from different countries (1999–2016)

|  | % International Collaboration | % National Collaboration | % Institutional Collaboration |
| --- | --- | --- | --- |
| Mexico | 37.9 | 16.8 | 36 |
| Argentina | 39.5 | 18.5 | 31.8 |
| Canada | 40.4 | 11.3 | 34 |
| France | 42.5 | 23.1 | 21.3 |
| Germany | 40.4 | 11.5 | 34.8 |
| Japan | 20.6 | 22.6 | 47.9 |
| Russian Federation | 28.1 | 14.6 | 40.2 |
| South Korea | 25.1 | 25.2 | 44.1 |
| Spain | 36.7 | 16.2 | 38.3 |
| United Kingdom | 38.2 | 13.1 | 26.9 |
| China | 15.9 | 24.9 | 54.7 |
| United States | 24.7 | 19.8 | 36 |

**Fig. 1.**
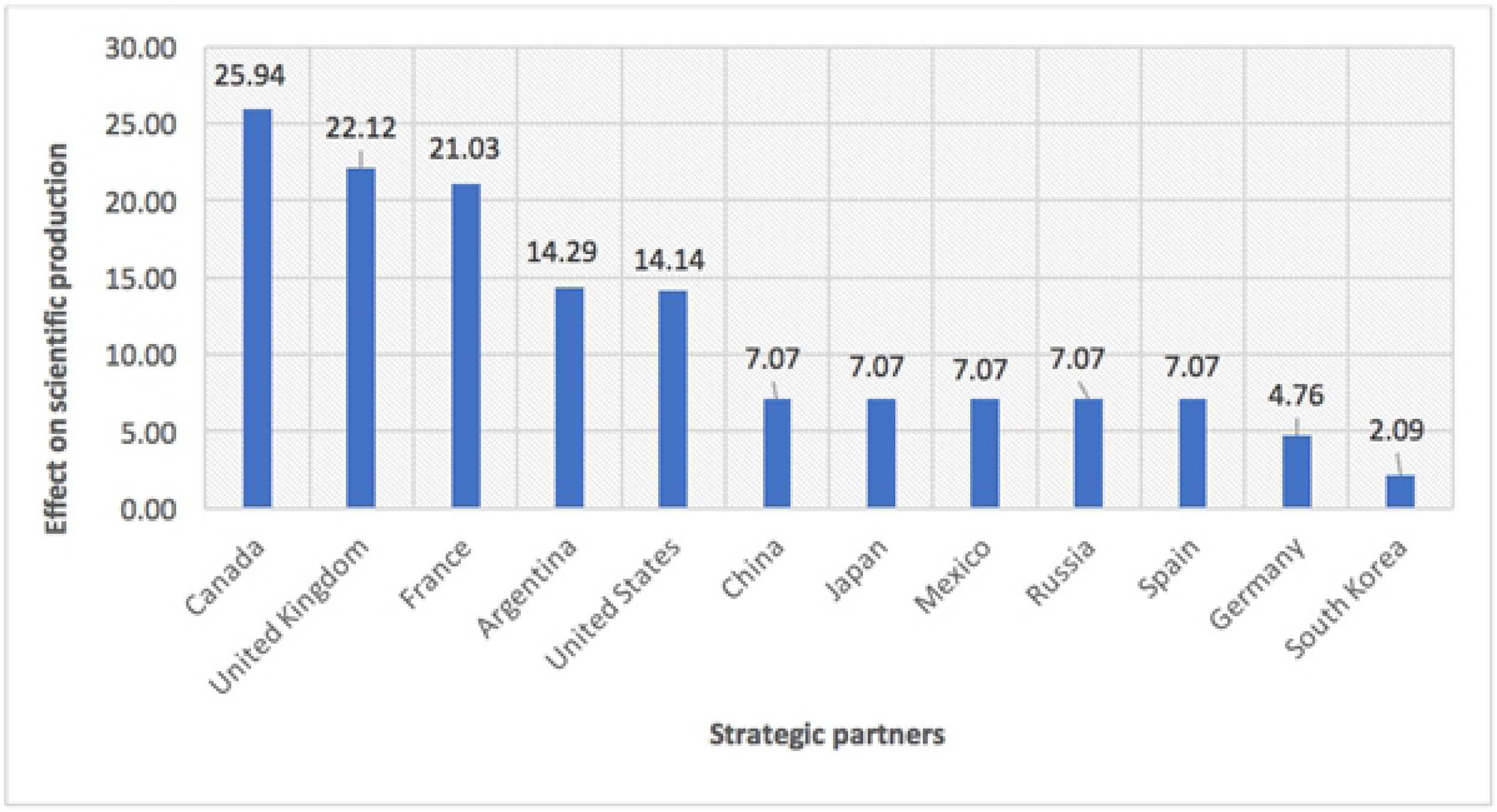
Effect on scientific production of different countries when variables GERD/GDP and A&RI tend to 0

When the economic investment and the presence of institutions tend to zero, the scientific production of Mexico is 7.068%. The dummy variables of countries that are not significant and therefore demonstrate behavior equal to that of Mexico include Japan, Russia, Spain, and China. The effect of these countries is the same as that of Mexico, i.e., when the GERD/GDP and A&RI variables tend to zero, the scientific production of these countries is 7.068%. When these variables tend to zero, the countries that surpass Mexico in percentage of scientific production are Argentina with production of 14.285%, France with 21.033%, and the United Kingdom with 22.123%. The maximum value comes from Canada with 25.936%.

As mentioned in the introduction, scientific production could be increased through collaboration. In the following table, we will check if these countries mentioned above have high levels of scientific collaboration.

Mexico has an international collaboration rate of 37.9%, and Argentina, France, the United Kingdom, and Canada have international collaboration rates of 39.5%, 42.5%, 38.2%, and 40.4%, respectively.

However, countries that behave like Mexico have a smaller percentage of documents in international collaboration and more in institutional collaboration. These countries are Japan, the Russian Federation, Spain, and China. Note that Mexico’s international collaboration (37.9%) and institutional collaborations (36%) are nearly the same. The only country with a lower value than the intercept is Germany, which has an effect of 4.756%, although it has a high percentage of international collaboration (40%). South Korea has a 2.087% lower value than Mexico and has lower international collaboration (25%) and more institutional collaboration (44%). International collaboration explains the highest values in production with respect to Mexico because these countries have a greater percentage of collaboration than Mexico.

Figure 2 shows the effect of the different countries on the scientific production relative to a 1% increase in GERD/GDP.

**Fig. 2.**
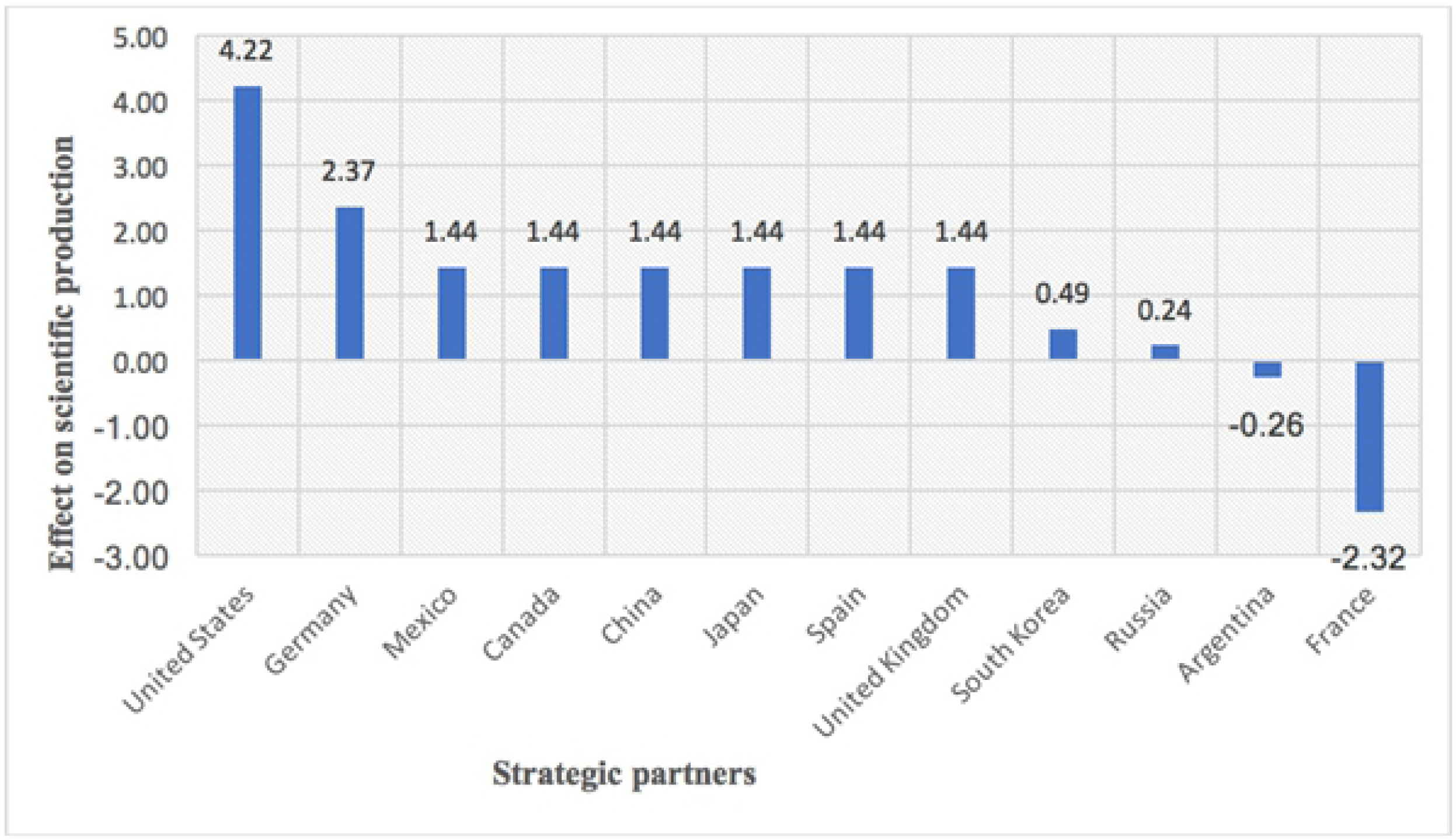
Effect on scientific production when GERD/GDP increases by 1% in all countries

If we increase the GERD/GDP of all countries by 1%, the scientific production increases in all of them, except for Argentina and France, whose scientific production diminishes and shows a negative effect. The fact is that Argentina had a fluctuating and low investment in GERD/GDP throughout 1996–2008. Despite the government’s low investment in science, scientific production continued to increase. In 2009, there was a boom in research investment, which was the maximum in the country’s history to date. Thus, from that time point to the present, investment has been increasing and has remained on the rise and stabilized since then. We show this fact more clearly in Figure 3. The data from 2009–2012 (GERD/GDP) with 2012/2015 (Scientific production) make the relationship significant in Argentina; however, as most data 1996- 2008 GERD/GDP have a negative relationship with respect to data from scientific production 1999-2011, the regression interprets it as negative because this is the majority. If this same study would be performed 10 years later and investment in Argentina continues to rise (as well as production), the sign would change and the relationship would be positive.

**Fig. 3.**
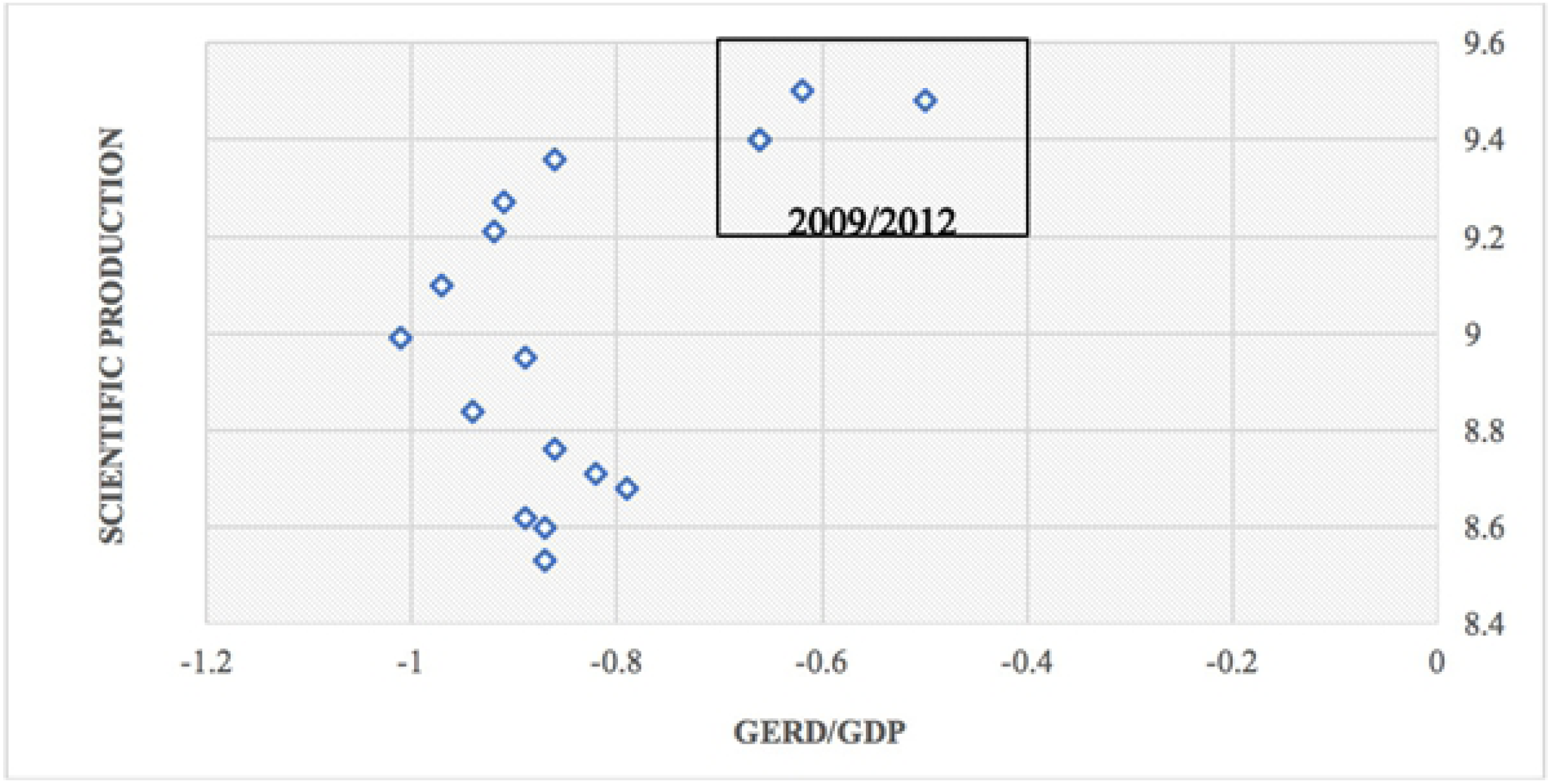
GERD/GDP versus scientific production of Argentina

In France, the trend is similar to that in Argentina. Investment in science during the years 1996–2008 was low, whereas production continued to rise in this period. In 2009, France shows the highest rise in research in 10 years. Figure 4 shows the relationship of GERD/GDP to scientific production. The most recent data show high values that make the slope positive. However, the estimation considers all values, which yields a negative relation.

**Fig. 4.**
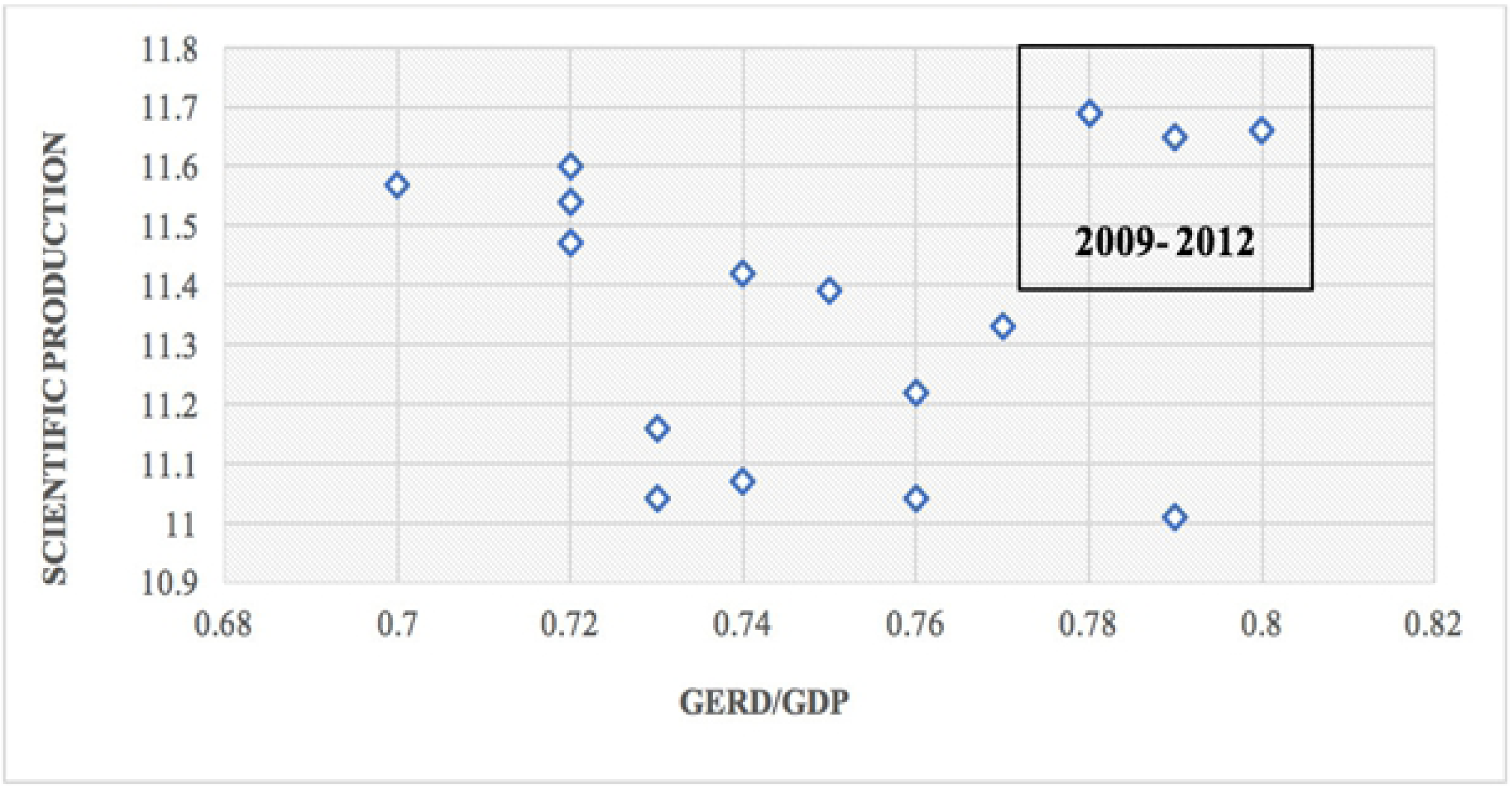
GERD/GDP versus scientific production of France

France is the fourth OECD (Organization for Economic Co-operation and Development) country behind the United States, Germany, and Japan relative to science investment. France also ranks sixth in the world in terms of the numbers of scientists.

If the GERD/GDP increases by 1%, the scientific production in the case of Mexico will increase by 1.44%. Countries that have the same effect as Mexico are Canada, China, Japan, Spain, and the United Kingdom. Countries that have a greater effect than Mexico are the United States and Germany. In contrast, countries that have a smaller effect than the reference country are Russia and South Korea.

Although Figures 3 and 4 show three values in the window 2009–2012, two of those values overlap and only one is shown.

**Fig. 5.**
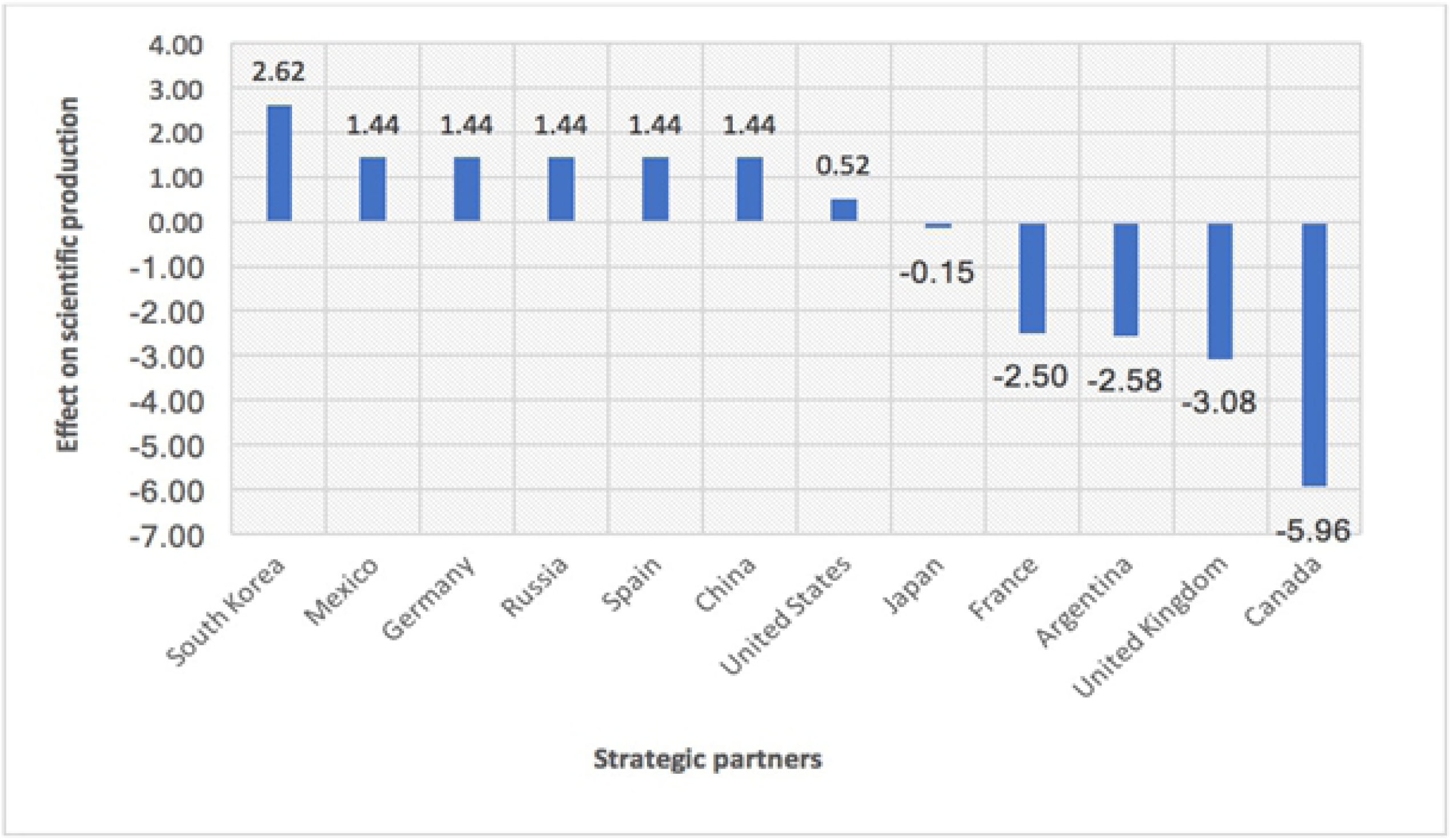
Effect on scientific production of different countries when A&RI increases by 1%

This figure reflects the fact that there are countries that tend to decentralize scientific production and others prefer to concentrate it among a lower number of institutions. Accordingly, if the number of institutions with at least 50% of Mexico’s scientific output increases by 1%, their production increases by 1.436%. Countries that behave like Mexico are Germany, Russia, Spain, and China.

In contrast, the United States shows an effect that is 0.523 less than Mexico. Note that the United States maintains the number of institutions as constant over the years; thus, the variable was excluded from regression. The countries that appear to concentrate their production among a smaller number of institutions over the years are Japan, France, Argentina, the United Kingdom, and Canada. In fact, scientific production decreases in these countries if the A&RI value increases by 1%. Here Japan decreases its scientific production by 0.152, France by 2.498, Argentina by 2.581, the United Kingdom by 3.083, and Canada by 5.959. It appears that the production of these countries is positioning them to have fewer and fewer institutions over time. These are the countries whose production would be higher if both the GERD/GDP and A&RI are reduced.

The countries with negative effects in A&RI are those that have effectively concentrated production into fewer institutions. In contrast, those with positive effects tend to diversify productivity across more institutions.

A forthcoming study will analyze the citations received by countries that concentrate scientific production among fewer institutions and those that decentralize production to more institutions. With this analysis, we expect to determine which of the two measures is more efficient.

Finally, we would like to compare our study with others previously done. Castellacci and Natera [52] also studied the evolution of innovation national systems using panel data. Unlike that, our study uses a shorter time window (17 years instead of 27), which reduces the probability of observing structural changes on national research policies that would affect its results. On the other hand, we permit heterogeneity in the intercept and coefficients of every country, whereas Castellacci and Natera [52] can only assess heterogeneity for a few groups of countries, either by Gross Domestic Product (GDP) or by continent.

## 4 Conclusions

We have presented a causal model of the scientific production of countries through multiple linear regression using panel data. Panel data simultaneously exploit temporal (17 years) and cross-sectional dimensions (countries); thus, including more observations the error is minimized. Therefore, the results are attributed to causality rather than correlation.

Although our initial hypothesis included five variables: GERD/GDP, Researchers (FTE), A&RI, subject areas, and total citations received influencing scientific productivity, only two variables complied with all statistical assumptions. These are: GERD/GDP and Academic and Research Institutions (A&RI) which are responsible for the 50% of the production.

We verified that the number of researchers, subject areas that comprise 50% of scientific production, and total citations would not satisfy the statistical assumptions. Therefore, we could not verify whether they have a causal effect on scientific production using multiple regression.

When multiple linear regression is performed without the country effect, an R^2^ value (coefficient of determination) of 0.73 is obtained, which means that these two variables explain the dependent variable by 73%. When the country effect is considered using panel data, the R^2^ value increases to 0.98 at a significance level of 0.05. With our panel data model we reduce the error from 0.602 to 0.125.

We could observe the difference in the behaviour of the countries with respect to Mexico, the reference country, through parameter estimation. With Mexico, the two independent variables are significant, which is not the case for all countries, such as China and Spain, whose coefficients were not significant, thereby indicating that they behave the same as the reference country (Mexico).

We obtained a predictive statistical model to explain scientific production. This model considers scenarios in which we assume increases to either of the two independent variables to determine its effect on scientific production (the dependent variable) and compare different effects.

The United States and Germany most effectively capitalize investment in research. For Argentina and France, it was not possible to demonstrate a positive effect of investment in production because we need to observe the phenomenon over a longer period.

Five of the countries analyzed tend to concentrate scientific production among only a few institutions. If the number of institutions that comprise 50% of the scientific production increases, then productivity will decrease in Japan, France, Argentina, the United Kingdom, and Canada. Note that the United States maintains a constant number of institutions over time.

The regression model will allow researchers to prognosticate future scientific production. This can be achieved in a way that, when we increase the GERD/GDP values, we can observe effects on scientific production. Similarly, we can also make forecasts relative to the A&RI variable. We believe that our causal multiple regression model can support the governments of each country be aware of the importance of increasing their investment on science and concentrating or diversifying research budget on institutions.

Finally, this paper will be relevant for public administrations, governments, private sectors, councils responsible for Science and Technology policies because they could make inferences about how through some increase in investment scientific production could be boosted in a period of time. This type of study can provide some insights for comparing the science and technology policies of a country with respect to those of a group of countries, in order to find what they do differently to improve their scientific productivity.

Furthermore, this is a general spectrum of 12 strategic partners. The same study could be done taking into account different countries. We encourage countries to create more scientific alliances, getting involved, committing their effort and experience to achieve a certain purpose in which they could benefit in a framework of common cooperation.

## Acknowledgments

We are grateful to Tecnologico de Monterrey, which supported this study through the Intelligent Systems Group. We also acknowledge the very valuable statistical guidance and assistance from Jose Enrique Montemayor Gallegos, Director of Academic Evaluation and Institutional Effectiveness at Tecnologico de Monterrey. His knowledge and expertise were fundamental in constructing the multiple regression model using panel data.

